# Lighting and circadian cues shape locomotor strategies for balance and navigation in larval zebrafish

**DOI:** 10.1101/2025.11.18.689084

**Authors:** Jiahuan Liu, Andy Kraja, Samantha N. Davis, Hannah Gelnaw, Naoroz Mahmood, David Schoppik, Yunlu Zhu

## Abstract

Most fish are inherently unstable and must swim to stabilize posture. How diurnal fish reduce activity at night while maintaining postural control remains unclear. We defined distinct locomotor strategies that larval zebrafish (*Danio rerio*) use to control posture and navigate the water column in response to light and circadian cues. In the dark, larvae maintain balance by swimming in long bouts with large nose-up rotations, compensating for nose-down drift accrued during prolonged inactivity. Effective postural compensation requires vestibular sensation from the utricle. By contrast, in the light, larvae navigate with short, frequent, and variable bouts. While lighting exerts a dominant, masking effect on the locomotor strategies, circadian rhythms modulate swim kinematics. Our results reveal distinct light-dark locomotor strategies and disentangle how ambient light and the internal clock jointly shape balance control and navigation. This work lays the foundation for understanding how external and internal cues interact to govern locomotor activity in freely moving diurnal animals.

## INTRODUCTION

Environmental lighting and the internal circadian clock interact to regulate most animals’ locomotor activity. The internal clock drives rhythmic behaviour [1, 2], while light acts both as a dominant zeitgeber that entrains the clock [3, 4] and as a direct modulator of behaviour through its masking effects [5–8]. Terrestrial animals that are active during the day rest at night by remaining still [9–11]. However, zebrafish (*Danio rerio*), like most fish species, lack a stable body plane [12, 13], requiring active swimming to maintain their preferred horizontal posture [14]. How diurnal fish stabilize posture despite reduced locomotor activity at night remains unknown.

Larval zebrafish are an ideal system for studying how lighting and circadian rhythms shape locomotor behaviour. They are more active during the day [15–22] and display light-induced masking of activity [23, 24]. Zebrafish larvae swim in discrete bouts separated by periods of inactivity [25, 26], making their locomotion easy to quantify. In the first few days post fertilization (dpf), larvae explore the water column to inflate their swim bladder at the surface [27, 28], hunt microorganisms [29], and adjust depth in response to illumination changes [23, 30, 31]. Zebrafish larvae experience a constant nose-down torque due to their front-heavy body plane [13, 14]. To maintain a preferred horizontal posture and navigate the water column effectively, they must sense and correct body tilts along the pitch (nose-up/nose-down) axis using vestibular feedback [14, 26, 32]. Although lighting and circadian cues influence activity levels in zebrafish larvae, their specific impact on the kinematics governing postural control and navigation remains unexplored.

We combined high-resolution behavioural recordings with photoperiodic manipulations to dissect how lighting and circadian rhythms shape locomotor strategies in larval zebrafish. We found that larvae adopt two distinct strategies depending on light availability and identified bout duration as a key kinematic parameter that classifies these two strategies. In the dark, larvae prioritized postural stabilization though long swim bouts accompanied by pronounced nose-up rotations. These rotations effectively compensated for nose-down postural drifts accrued during prolonged inactivity. In the light, larvae adopted frequent, short swim bouts with high directional variability, facilitating horizontal exploration. Transitions between lighting conditions induced quick strategy switching. Interestingly, while lighting exerts a dominant masking effect on locomotor strategies, circadian rhythms modulate swim kinematics. Postural control and navigation exhibited rhythmic fluctuations in constant darkness. In conclusion, our study disentangles the contributions of lighting and circadian rhythms to locomotor activity, revealing how a simple vertebrate adjusts locomotor strategies to balance and navigate in response to external and internal cues.

## MATERIALS AND METHODS

### Fish husbandry

All procedures involving larval zebrafish (*Danio rerio*) have been approved by the Institutional Animal Care and Use Committee (IACUC) at New York University Langone Health. Adult zebrafish were maintained at 28.5 °C under a standard 14/10-hour light/dark cycle with lights on from 9 a.m. to 11 p.m. Fertilized embryos were derived from in-crosses of wild-type zebrafish with a mix of AB/WIK/TU/SAT/NHGRI-1 backgrounds. During the first day after birth, embryos were raised at densities ranging from 20 to 50 in 10 cm petri dishes, each containing 25 to 40 mL of E3 medium with 0.5 ppm methylene blue. At 1 day post-fertilization (dpf), larvae were transferred to E3 medium without methylene blue. Starting from 5 dpf, larvae were raised at a density around 20 per dish and fed with cultured rotifers (Reed Mariculture). Larvae were kept in incubators set to a 28.5 °C 14/10-hour light/dark cycle until behavioural assessment.

### Behavioural measurements

We used the Scalable Apparatus to Measure Posture and Locomotion (SAMPL) to record behaviour of freely swimming zebrafish larvae [25]. The apparatus, experimental procedure, program, and analysis pipeline have been detailed previously [25]. Each behaviour box was equipped with a cool white LED strip to control photoperiodic treatments, illuminating the chamber with cool white light measured approximately between 50 to 150 lux with irradiance between 0.61 to 3.91 W/m^2^. Constant light (LL) and constant dark (DD) conditions were generated by keeping the white LED on or off, respectively. For the light/dark (LD) condition, the white LED followed a standard 14/10-hour light/dark cycle with the light on from 9 a.m. to 11 p.m.

For the 2-day recordings, 5 to 7 larvae were transferred into each custom-designed behaviour chamber filled with 30 mL of E3 medium at 7 dpf. The water filled a portion of the chamber that measured approximately 50 mm (L) × 50 mm (H) × 13 mm (W). Each chamber was positioned between a camera and a 940 nm infrared light source inside a light-tight box. Behavioural recordings began before 11 a.m. on the first day and lasted for approximately 48 hours. The recording field of view was a 20 mm × 20 mm square in the centre of the lower half of the chamber. We recorded the coordinates and angle along the pitch axis (nose-up/nose-down) of larvae in the field of view in real time at 166 Hz. After approximately 24 hours, recordings were paused for 30 minutes, where each chamber was fed with 1 to 2 mL of cultured rotifers. Behaviour data were collected from 105 and 98 fish under dark and light conditions, respectively, across five experimental repeats. Each experimental repeat included data collected from larvae of the same clutch (siblings). For each condition in each repeat, data were collected from three SAMPL boxes containing approximately 20 fish in total. Additional information on sample sizes is provided in the figure legends and tables.

For the 6-day recordings used for circadian analyses, each chamber contained only a single larva. Larvae raised in incubators under normal light-dark cycles were transferred to behaviour chambers at 4 dpf together with 3 mL of rotifer culture per chamber. Each chamber was covered with a piece of Parafilm punctured with small holes to reduce water evaporation. Recordings were performed for 6 days without additional feeding or opening of the boxes. This dataset was collected from 18, 19, and 17 fish under DD, LD, and LL conditions, respectively, across 6 experimental repeats. Additional information on sample size is provided in the figure legends and tables.

### Lesions of lateral line hair cells

At 7 dpf, larvae were treated with 10 µM copper sulfate (CuSO_4_) in E3 medium for 90 minutes. Control siblings were handled identically and transferred to E3. After the treatment, larvae were washed in E3 and transferred to behavioural chambers. Following 24 hours of behavioural assessment, the CuSO_4_ treatment was repeated to prevent hair cell regeneration. Quantification of lateral line hair cell loss following the CuSO_4_ treatment can be found in [33].

### Behavioural analysis

Raw behavioural data were preprocessed using pipelines previously published with the SAMPL apparatus [25]. First, swim speed was calculated from recorded coordinates. A speed threshold of 5 mm/s was used to detect swim bouts. Then, we segmented the time-series data into swim bouts and inter-bout intervals (IBIs) by extracting a 450 ms window around the time of peak speed for each bout. Swim bouts were then aligned at the time of peak speed for subsequent analyses.

Next, we analysed the preprocessed data to extract kinematic parameters. For each bout, we calculated the peak speed, bout duration, swim displacement, direction, and body rotation. To capture the temporal dynamics of swim speed, we defined the bout duration as the duration of time that swim speed remained above 50% of its peak, i.e. the width at half maximum of the speed profile. Swim displacement was determined as the Euclidean distance travelled during the period when speed remained faster than 5 mm/s. Swim direction was defined as the instantaneous trajectory at the time of the peak speed on the pitch (nose-up/nose-down) axis. To assess rotation, we analysed the body angle along the pitch axis relative to horizontal. Bout rotation was defined as the net change in pitch angle between –250 ms and +200 ms relative to the time of peak speed. IBI postural drift was defined as the pitch difference between the end of one bout and the start of the next, representing the passive postural drift between consecutive swim bouts. Postural compensation residual was calculated by the addition of IBI postural drift and the subsequent bout rotation. Residual variability was defined as the median absolute deviation of the compensation residual.

Navigation metrics were computed from epochs containing multiple swim bouts. Displacement per second was calculated using epochs comprising five consecutive bouts, defined as the Euclidean distance between the starting positions of the first and fifth bouts, divided by the time elapsed between them. Directional change was defined as the absolute change in swim direction between two consecutive bouts [26]. The absolute directional changes between bouts 1 and 5 were summed and divided by elapsed time to obtain the directional change per second.

To visualize circadian effects, we extracted data from 9 a.m. on the second day of behaviour (5 dpf) to 9 a.m. on the sixth day (9 dpf). Kinematic measurements were binned into 1-hour intervals and summarized as the median value for each fish. For circadian analyses, the binned values were pooled and detrended by subtracting the fitted linear drift estimated by linear regression. Lomb-Scargle periodograms were then applied to the detrended values, with the peak period searched over 12 to 36 hours. The dominant period identified for each condition was used for subsequent Cosinor fitting.

### Statistical analysis

For each behavioural parameter, we assessed its distribution to select the appropriate statistical tests. None of the parameters followed a normal distribution, so we proceeded with non-parametric methods for all analyses.

In summary tables, results from all experimental repeats were pooled, and parameter distributions were reported as medians with interquartile ranges (IQR). P-values were calculated using the median test. Effect sizes for median tests were estimated as 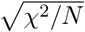, a transformation commonly used to express chisquared statistics as a standardized effect size analogous to correlation coefficients [34]. Scott’s rule was applied to determine the number of bins for all histograms [35].

For comparisons reported in the main manuscript, each experimental repeat was summarised by its median, and the means of these medians were compared across conditions using independent *t*-tests. *P*-values and Cohen’s *d* were reported in the main text and figure legends.

Correlations of bout rotation and IBI drift were examined using bi-square regression, a robust method that reduces sensitivity to extreme values. Statistical analyses of correlation coefficients were performed using two-way ANOVA with post-hoc Tukey HSD tests with the Bonferroni correction applied to control for Type I errors.

Statistical analyses of LombScargle periodograms were performed using a permutation test. The per-fish 1 h median values were randomly reassigned to the original time labels 1,000 times to generate a null distribution of periodogram power. P values were calculated as the proportion of permutations yielding a maximum periodogram power equal to or greater than that observed in the original dataset.

### Data and code availability

All raw data and code for analysis have been uploaded to the Open Science Framework: 10.17605/OSF.IO/36TFU.

## RESULTS

### Larvae employ distinct swimming kinematics under dark and light

We first examined how lighting affects kinematics of freely swimming zebrafish larvae. We recorded larval activity using a custom-designed Scalable Apparatus to Measure Posture and Locomotion (SAMPL) [25]. SAMPL records larvae from their side (Figure 1A) and measures swim speed and posture on the pitch (nose-up/nose-down) axis (Figure 1B) at a frame rate of 166 Hz [25]. Zebrafish larvae swim in short, discrete bouts interspersed with inactive periods called inter-bout intervals (IBI) (Figure 1B). Due to their anteriorly positioned centre of gravity relative to the centre of buoyancy (Figure 1C), larvae are inherently unstable and rotate nose-down during IBIs [14](Figures 1B and 1C). We recorded swim parameters from 7 to 9 days post-fertilization (dpf) and extracted a 450 ms window of activity for each bout (Figure 1D). We generated light-dark (LD) and dark-dark (DD) photoperiodic conditions by turning the daylight LED in the apparatus on or off during the day time, respectively, and compiled day-time data for analysis (Figure 1E).

**Figure 1:**
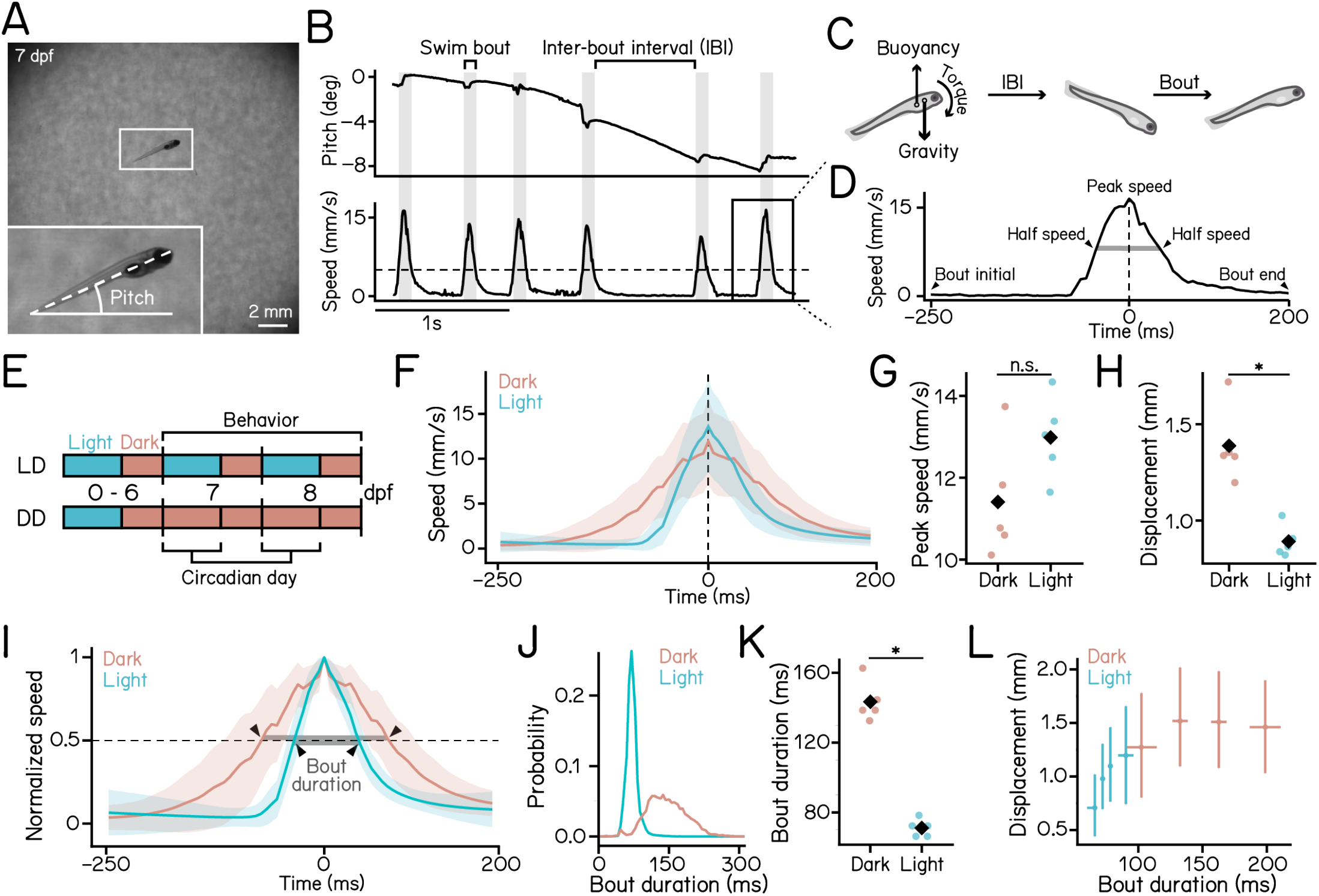
Measuring posture and locomotion in freely swimming zebrafish larvae reveals distinct kine-matics in dark and light. **(A)** Representative image of a 7 dpf zebrafish larva recorded in the SAMPL apparatus. Pitch angle is defined as the angle of the trunk axis (dashed line) relative to the horizontal. Positive values indicate nose-up posture. **(B)** Example epoch containing multiple swim bouts, with pitch angle (top) and swim speed (bottom) plotted over time. The horizontal dashed line marks the 5 mm/s threshold used for bout detection. Grey vertical bars indicate the duration of swim bouts. The inter-bout interval (IBI) is defined as the time between two bouts. **(C)** Schematic illustrating the forces acting on a larva along the vertical axis. Because the centre of mass lies anterior to the centre of buoyancy, larvae experience a nose-down rotation during IBIs and correct their posture through swim bouts. **(D)** Time series of swim speed during a swim bout. The vertical dashed line indicates the time of peak speed. Bout duration (grey bar) is defined as the duration between the two half-speed points (arrowheads). **(E)** Schematic illustration of experimental paradigm. Larvae were raised under a standard 14–10 light-dark cycle from 0 to 6 dpf and transferred into the SAMPL apparatus for behavioural recording from 7 to 9 dpf under either light-dark (LD) cycle or constant-dark (DD) conditions. Data from the zeitgeber day were used for analysis. **(F)** Swim speed plotted as a function of time under dark (red) and light (cyan) conditions. Solid lines represent the mean across 5 experimental repeats; shaded areas indicate ±1 standard deviation (SD) across repeats. **(G)** Peak swim speed under dark (red) and light (cyan). Median values for each experimental repeat are plotted as dots. Black diamonds indicate group means (p = 8.053e-02, Cohen’s d = 1.265, t-test). **(H)** Bout displacement under dark (red) and light (cyan). Median values for each experimental repeat are plotted as dots. Black diamonds indicate group means (p = 7.831e-04, Cohen’s d = 3.314, t-test). **(I)** Time series of normalized swim speed. For each bout, speed is normalized to its peak value. Solid lines represent the mean across 5 experimental repeats; shaded areas indicate ±1 SD. Bout duration is defined as the duration (grey bar) between the two time points where speed is at 50% of the peak value (arrowheads). **(J)** Histogram of bout duration under dark (red) and light (cyan) conditions. **(K)** bout duration under dark (red) and light (cyan) conditions. Median values for each experimental repeat are plotted as dots. Black diamonds indicate group means (p = 1.315e-06, Cohen’s d = 8.090, t-test). **(L)** Swim bout displacement plotted as a function of bout duration. For each condition, data were grouped into four equal-sized quartiles based on bout duration. Crossing points indicate the median of displacement and bout duration of each quartile. Horizontal and vertical error bars represent the interquartile ranges (IQRs). n = 23875/98722 bouts from 105/98 fish for dark/light over 5 experimental repeats. See also Table 1.

Compared to siblings under light conditions, larvae in the dark reached a comparable peak speed (Figures 1F and 1G; dark: 11.409 ± 1.445 mm/s vs. light: 12.982 ± 1.003 mm/s, *p* = 8.053e-02) and exhibited greater swim displacement (Figure 1H; dark: 1.388 ± 0.195 mm vs. light: 0.891 ± 0.082 mm, *p* = 7.831e-04). The difference in displacement was marginally significant, whereas peak speed did not differ significantly between groups. The relatively small number of experimental repeats limited statistical power. We therefore validated these findings by pooling swim bouts within each condition and comparing at the individual bout level (Table 1). This analysis indicated that larvae in the dark swim at lower peak speed but cover greater distances.

**Table 1:**
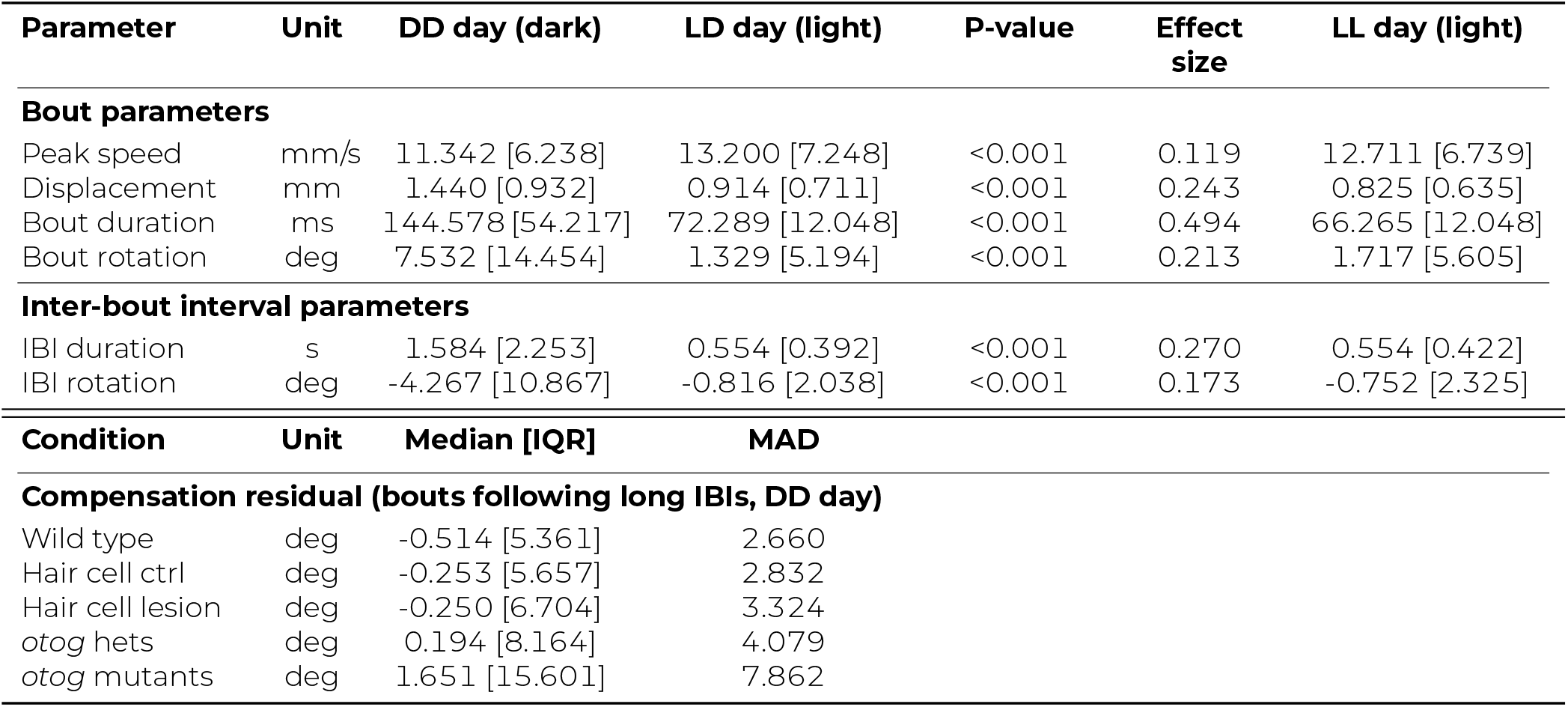
Kinematic parameters under DD, LD, and LL during day time. Refer to Figures 1–2. Data from 5 experimental repeats were pooled to calculate median [IQR]. n = 23875/98722/100674 day-time bouts from 105/98/98 fish for wild type DD/LD/LL. Median test p-values reported for DD day vs. LD day comparisons; Effect size estimated using standardized chi-squared statistics. n = 8336/2994 bouts from 114/114 fish for hair cell control/lesions. N = 3106/1716 bouts from 99/136 fish for *otog* hets/mutants. MAD: median absolute deviation. See Methods for details.

Notably, we found that dark bouts exhibited more gradual changes in speed (Figure 1F). To quantify the dynamics of the speed profile, we normalized each swim bout by its peak speed and measured the bout duration, defined as the duration during which larvae moved at speeds greater than half their peak speed (Figure 1I). Larvae in dark showed significantly longer bout durations compared to their siblings in light (Figures 1J and 1K; dark: 143.373 ± 11.588 ms vs. light: 71.084 ± 5.040 ms, *p* = 1.315e-06).

These findings led us to hypothesize that larvae achieve greater displacement through longer bouts. We plotted displacement as a function of bout duration and observed a positive correlation in the light condition (Figure 1L, cyan), indicating that increased bout duration is associated with greater displacement. However, this relationship was absent in bouts recorded in the dark (Figure 1L, red), suggesting that larvae employ a different locomotor regime without light, and leaving open the question of the functional significance of extending bout duration in the dark.

These results demonstrate that zebrafish larvae exhibit distinct swim kinematics in light versus dark conditions, and that *bout duration* reliably differentiates activity between these conditions.

### Swim bouts in the dark stabilize posture

Zebrafish larvae rotate in the pitch axis during swim bouts [14, 25, 26]. Longer bout durations allow more time for rotation. Given the significantly longer bout durations in the dark, we hypothesized that larvae use these extended swim bouts to achieve greater body rotation.

We measured rotations during swim bouts (Figure 2A) and observed significantly greater nose-up rotations in dark bouts (Figures 2B and 2C; dark: 5.724 ± 3.042°, light: 1.053 ± 0.504°, *p* = 9.538e-03). Notably, bout rotations were positively correlated with bout duration in the dark (Figure 2D, red), indicating that larvae extend their bout duration to achieve greater nose-up rotations.

**Figure 2:**
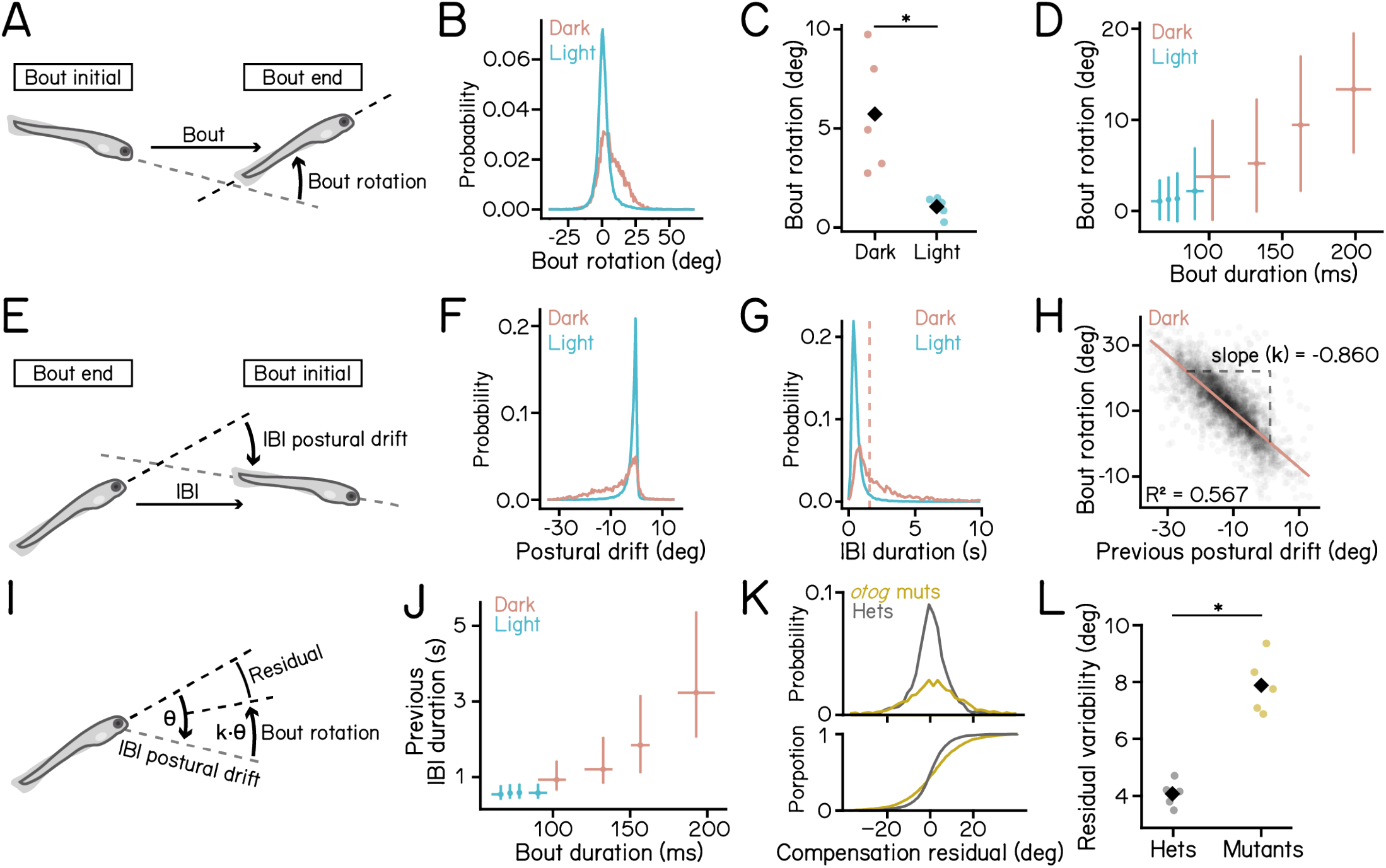
Swim strategy in dark compensates for postural drifts accrued during prolonged inactivity. **(A)** Schematic illustration of bout rotations, defined as the change in pitch angle during a swim bout. **(B)** Histogram of bout rotation under dark (red) and light (cyan) conditions. **(C)** Bout rotation under dark (red) and light (cyan) conditions. Medians for each experimental repeat are plotted as dots. Black diamonds indicate group means (p = 9.538e-03, Cohen’s d = 2.142, t-test). **(D)** Bout rotation plotted as a function of bout duration. For each condition, data were divided into four equalsized quartiles based on bout duration. Crossing points indicate the median rotation and bout duration of each quartile. The horizontal and vertical error bars represent IQRs. **(E)** Schematic illustration of IBI postural drift, defined as the change in the pitch angle from the end of the previous bout to the beginning of the current bout. **(F)** Histogram of postural drift under dark (red) and light (cyan) conditions. **(G)** Histogram of postural drift under dark (red) and light (cyan) conditions. The dashed red line indicates the median of IBI duration in the dark. Bouts with IBIs longer than this median are classified as long IBIs. **(H)** Scatter plot of bout rotation following long IBIs vs. previous IBI postural drift in dark. The red line represents the robust bi-square regression fit (slope k = −0.860, coefficient of determination *R*^2^ = 0.567). **(I)** Schematic illustration of compensation for postural drift following long IBIs. Larvae rotate nose-up proportionally to the amount of nose-down drift during the previous IBI. Compensation residual is defined as the sum of IBI postural drift and Bout rotation. **(J)** Previous IBI duration plotted as a function of bout duration. For each condition, data were divided into four equal-sized quartiles based on bout duration. The horizontal and vertical error bars represent IQRs. Crossing points indicate median of duration and bout duration of each quartile. **(K)** Histogram (upper) and cumulative distribution (lower) of compensation residuals of *otog*^-/-^ mutants (yellow) and heterozygous control (grey). Results from bouts in the dark with long proceeding IBIs were plotted. **(L)** Residual variability of *otog*^-/-^ mutants (yellow) and heterozygous control (grey). Medians for each experimental repeat are plotted as dots. Black diamonds indicate group means (p = 5.652e-05, Cohen’s d = 4.881, t-test). n = 23875/98722 bouts from 105/98 fish for dark/light over 5 experimental repeats. n = 3143/34285 middle bouts from 3-bout sequences were selected for calculation of IBI postural drifts for dark/light conditions. For the *otog* mutant dataset, n = 3106/1716 bouts with long preceding IBIs from 99/136 fish for hets./mutant over 5 experimental repeats. See also Figure S1 and tables 1 and 2.

Why do they rotate more nose-up in the dark? Because larvae experience a constant nose-down torque, we reasoned that in the dark they would accumulate larger nose-down postural drifts during inter-bout intervals (Figure 1C). We quantified kinematics during IBIs (Figure 2E) and found that larvae in the dark showed increased nose-down drifts (Figure 2F) and prolonged IBI durations (Figure 2G). These results support the hypothesis that larvae rotate nose-up to compensate for nose-down drifts accrued during prolonged IBIs in the dark.

**Table 2:** Correlation of bout rotation and IBI rotation under dark and light. Refer to Figure 2. Wild-type data from 5 experimental repeats were pooled and analysed using bi-square regression. Middle bouts from 3-bout sequences were selected for analysis. n = 3143/34285 day-time bouts from 105/98 fish for DD/LD over 5 experimental repeats. The hair-cell lesion dataset contains 8336/2994 day-time bouts from 114/114 fish for control/lesions over 5 experimental repeats.

| IBI length | Relation | DD day (dark) |  |  | LD day (light) |  |  |
| --- | --- | --- | --- | --- | --- | --- | --- |
| | | Slope | $R^2$ | Bootstrapped p-value | Slope | $R^2$ | Bootstrapped p-value |
| Wild type |  |  |  |  |  |  |  |
| Long | Previous IBI | -0.860 | 0.567 | <0.001 | -0.360 | 0.164 | <0.001 |
|  | Next IBI | -0.874 | 0.172 | <0.001 | -0.648 | 0.088 | <0.001 |
| Short | Previous IBI | -0.986 | 0.185 | <0.001 | -0.753 | 0.034 | <0.001 |
|  | Next IBI | -0.712 | -0.093 | <0.001 | -0.282 | 0.011 | <0.001 |
| Lateral-line hair cell lesion control |  |  |  |  |  |  |  |
| Long | Previous IBI | -0.852 | 0.451 | <0.001 |  |  |  |
| Short | Previous IBI | -1.131 | 0.226 | <0.001 |  |  |  |
| Lateral-line hair cell lesioned |  |  |  |  |  |  |  |
| Long | Previous IBI | -0.756 | 0.545 | <0.001 |  |  |  |
| Short | Previous IBI | -1.004 | 0.371 | <0.001 |  |  |  |

To directly test this hypothesis, we examined correlations between IBI postural drift and subsequent bout rotation. We categorized swim bouts into those with long vs. short IBIs based on the median dark IBI duration (Figure 2G, dashed vertical line). We plotted bout rotations against IBI postural drifts during either the preceding or following IBI (Figure S1). Bout rotations in the dark were negatively correlated with postural drift during long IBIs, whereas no such correlation was observed under the light condition (Figures S1A to S1C). Interestingly, bout rotations were strongly linearly correlated with drift during the preceding IBIs (Figure 2H, *R*^2^ = 0.567), but not the following IBIs (Figure S1B, *R*^2^ = 0.172, quantified in Figure S1D), suggesting that larvae compensate for prior postural drift rather than anticipate future drift. To measure the degree of compensation, we defined the absolute slope of the best-fit line as the compensation gain (Figure 2H). Swim bouts in the dark exhibited a gain of 0.860(Figure 2H), indicating that following long IBIs larvae effectively compensate for nose-down rotations accrued during the IBI through swim bouts (Figure 2I). Correspondingly, bouts after longer IBIs in the dark exhibited longer bout durations (Figure 2J). These results demonstrate that, larvae perform nose-up counter rotations to compensate for nose-down drifts accrued during long periods of inactivity.

Building upon this observation, we further examined whether the IBI duration or the magnitude of postural drift determines the amount of compensatory rotation. Fish rely on the lateral line to sense water flow and the water-air interface [28, 36]. Lateral line sensation is critical for many locomotor behaviours including predator detection, prey, schooling, and rheotaxis [37–42]. Recent work show that ablation of lateral-line hair cells after swim bladder inflation in larval zebrafish reduces the size of the swim bladder and makes them more negatively buoyant [33]. To dissociate IBI duration with the amount of passive postural drift, we took the advantage of a this balance challenge that exacerbates the nose-down destabilizing torque. We used low-concentration copper sulfate to ablate the lateral-line hair cells and observed increased nose-down angular velocity during inactivity (Figure S1E). Hair-cell-lesioned larvae showed greater nose-down rotation during IBI despite having shorter IBI durations (Figures S1F and S1G), and rotated more nose-up during swim bouts (Figure S1H). Excitingly, we observed a similar linear relationship between bout rotation and postural drift during preceding IBIs (Figure S1I and table 2, *R*^2^ = 0.545) and measured a compensation gain of 0.756. The robust compensation for postural drift following lateral-line hair cell ablation demonstrate that this specific behaviour requires little lateral-line sensory input. Thus, we conclude that the amount of compensatory rotation is not determined by the duration of the inactive period but instead by the amount of passive postural drift accrued.

Next, we asked whether the sensation of gravity contributes to the compensatory rotation. We used the *otogelin* mutant [43], which lacks the utricular otolith for the first two weeks of life (Figure S1J) and thus exhibits impaired gravity sensation [44–47]. We selected mutant larvae with successfully inflated swim bladder and examined their swim bouts following long IBIs. To assess the effectiveness of compensatory rotation, we quantified the compensation residual, defined as the sum of IBI postural drift and the bout rotation (Figure 2I). Mutants showed a broader distribution of compensation residual than their heterozygous siblings (Figure 2K), reflected in significant higher residual variability in mutants (Figure 2L; heterozygous siblings: 4.070 ± 0.458°, mutants: 7.892 ± 1.008°, *p* = 5.652e-05). These results show that vestibular sensation via the inner-ear otolith is essential for effective postural compensation.

We conclude that larvae in the dark use long swim bouts with pronounced nose-up rotations to compensate for nose-down postural drifts during prolonged inactivity, reflecting a locomotor strategy to stabilize posture.

### Locomotor strategy in the light favours exploration

Previous studies reported that light increases activity level and swim distance of larval zebrafish [19, 48], thereby promoting effective exploration [49]. In contrast, we found that although larvae initiated swim bouts more frequently in the light (Figure 2G), individual bouts generated shorter displacements than those in the dark (Figure 2D). This discrepancy prompted us to investigate how larvae chained individual swim bouts to explore the water column effectively in the light.

We compared swim trajectories between light and dark conditions and observed more swim bouts and overall longer distance traveled in the light (Figure 3A). To quantify swim displacement efficiency, we calculated displacement per second by dividing the Euclidean displacement of the bout sequence by the total duration (refer to Methods). We found that larvae in the light travelled significantly farther than those in the dark over the same period (Figures 3B and 3C; dark: 2.317 ± 0.831 mm/s, light: 3.980 ± 0.556 mm/s, *p* = 5.882e-03). Larvae primarily swim horizontally in the light while performing more climbs in the dark (Figure 3D). These results demonstrate that larvae travel farther in the light by combining short individual swim bouts with faster swim rates.

**Figure 3:**
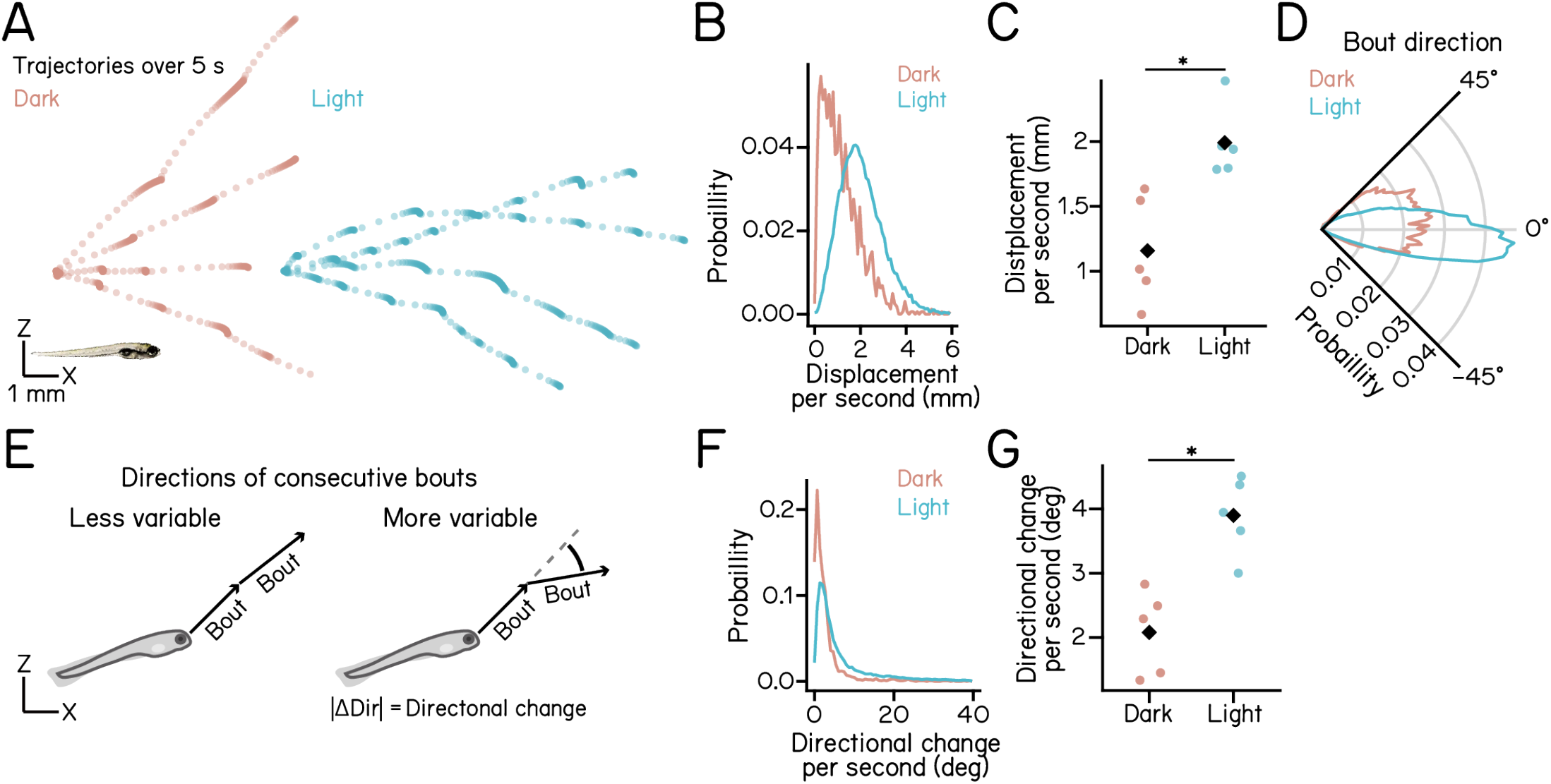
Swim bouts in the light travel further with greater directional variability per unit time. **(A)** Five-second trajectories of larvae under dark (red) and light (cyan) conditions. Four trajectories moving left to right are shown per condition. Dots indicate fish positions at 30 ms intervals. A 7 dpf larva is shown at the bottom left for scale. **(B)** Histogram of Euclidean displacement per second under dark (red) and light (cyan) conditions. **(C)** Per-second Euclidean displacement under dark (red) and light (cyan) conditions. Medians for each experimental repeat are plotted as dots. Black diamonds indicate group means (p = 5.882e-03, Cohen’s d = 2.352, t-test). **(D)** Polar histogram showing distributions of bout directions. **(E)** Schematics illustrating directional changes across bouts, defined as the sum of absolute changes in swim direction between consecutive bouts. Arrows indicate swim bout directions. **(F)** Histogram of directional change per second under dark (red) and light (cyan) conditions. **(G)** Per-second directional change under dark (red) and light (cyan) conditions. Medians for each experimental repeat are plotted as dots. Black diamonds indicate group means (p = 1.862e-03, Cohen’s d = 2.881, t-test). n = 7980/120010 sets of consecutive bouts from 105/98 fish for dark/light over 5 experimental repeats. n = 23875/98722 swim bouts for (D). See also Table 3.

The ability to change movement direction is essential for environmental exploration. Previously, we measured changes in swim directions on the vertical axis across consecutive bouts (Figure 3E) and found that larvae in the dark swim with stable trajectories, enabling effective climbs and dives [26]. Here, we examined how short, frequent swim bouts in the light influence changes of swim directions.

**Table 3:** Swim displacement and directional changes of larvae during day time of the recording. Refer to Figure 3. Data from 5 experimental repeats were pooled to calculate median [IQR]. P-values for DD vs. LD comparisons are from the median test. n = 23875/98722/100674 day-time bouts from 105/98/98 fish for DD/LD/LL for individual bout displacement. n = 7980/120010/132090 sets of 5 consecutive bouts from 105/98/98 fish for DD/LD/LL for average displacement and directional change calculation. Median test p-values reported for DD day vs. LD day comparisons; Effect size estimated using standardized chi-squared statistics. See Methods for details.

| Parameter | Unit | DD day (dark) | LD day (light) | P-value | Effect size | LL day (light) |
| --- | --- | --- | --- | --- | --- | --- |
| Displacement per second | mm | 1.019 [1.117] | 1.977 [1.258] | 1.067e-154 | 0.166 | 1.897 [1.270] |
| Bout displacement | mm | 1.440 [0.932] | 0.914 [0.711] | <0.001 | 0.243 | 0.825 [0.635] |
| Bout direction | deg | 12.971 [30.907] | -0.095 [19.875] | <0.001 | 0.183 | 0.828 [20.144] |
| Directional change per second | deg | 1.959 [2.391] | 4.005 [6.197] | 1.318e-129 | 0.151 | 3.800 [5.775] |
| Directional change between bouts | deg | 4.809 [8.064] | 3.792 [7.043] | 4.199e-49 | 0.041 | 3.648 [6.552] |

We first calculated the average change in direction between adjacent bouts. Bouts in the light showed slightly smaller directional changes than those in the dark (dark: 5.541 ± 0.701°, light: 4.123 ± 0.668°, *p* = 1.129e-02). We then assessed how directional changes accumulate over time. We calculated the rate of directional changes by dividing the total magnitude of change across a bout sequence by its duration (refer to Methods). Interestingly, larvae in the light exhibited significantly greater directional changes per unit time than those in the dark (Figures 3F and 3G; dark: 2.080 ± 1.315°/s, light: 3.901 ± 1.211°/s, *p* = 1.862e-03). This apparent discrepancy is explained by the shorter bout intervals in the light (Figure 2G), which increase bout frequency and thereby accelerate the accumulation of directional change over time.

These results led us to conclude that, under light conditions, larvae employ short, frequent bouts to achieve greater displacement and higher directional variability per unit time, reflecting a strategy that favours exploration.

### Dissociation of circadian and lighting effects on locomotion

The circadian clock internally regulates locomotor activity. We hypothesized that kinematics of postural control and navigation are rhythmically modulated by the circadian clock.

To capture circadian effects, we adapted a previously established paradigm [18], where embryos and larvae were exposed to LD cycles for 4 days and then transferred to the behavioural apparatus under LD, DD, and constant light (LL) conditions (Figure 4A). Larvae received a single dose of rotifers at 4 dpf. We then examined their behaviour over 5 days from 5 to 9 dpf (Figure 4A).

**Figure 4:**
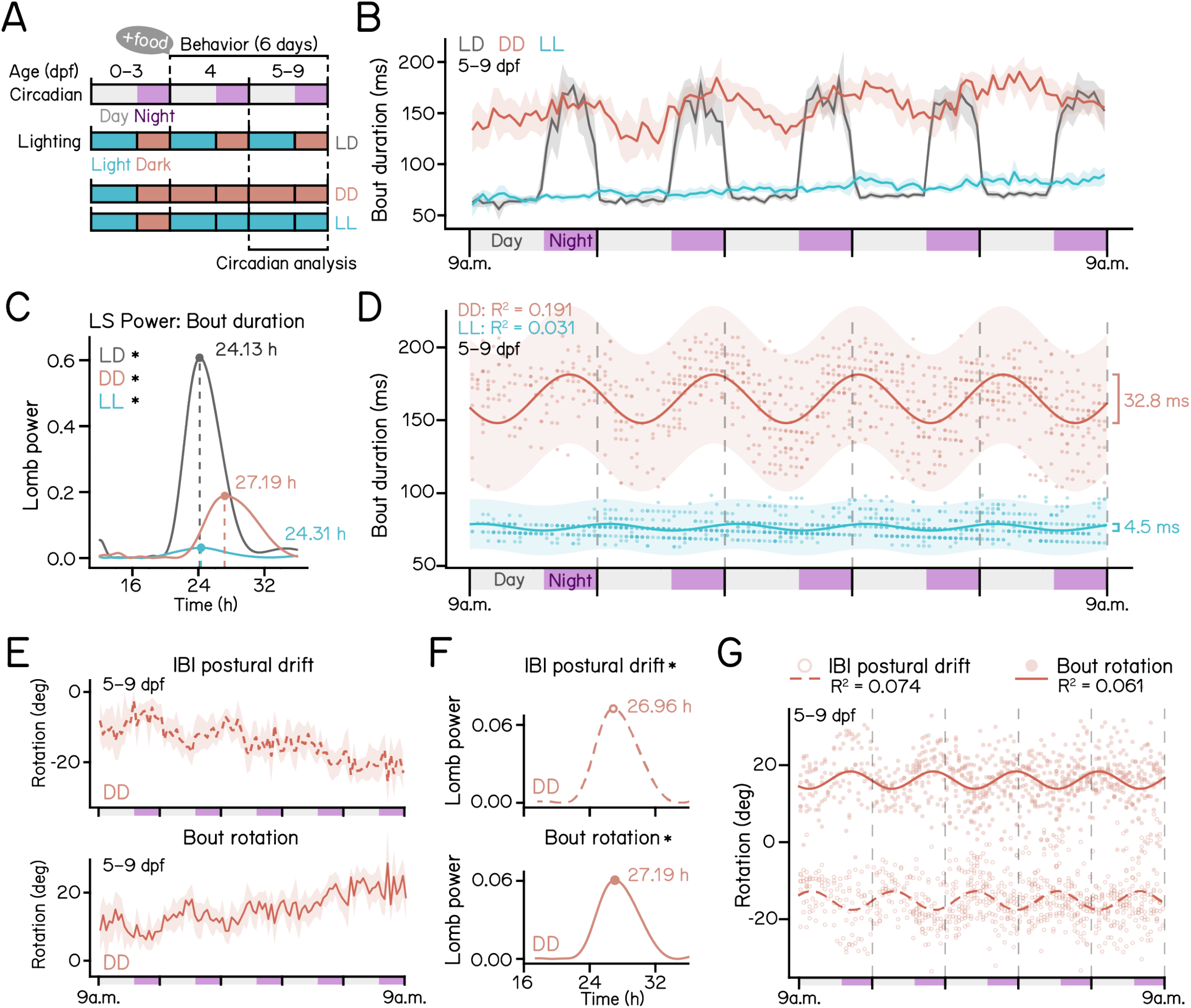
Dissociation of lighting and circadian effects on swim kinematics. **(A)** Experimental paradigm for circadian analysis. Data from 5 to 9 dpf were used for analysing circadian effect. **(B)** Bout duration plotted as a function of time under three lighting conditions: DD (red), LD (grey), and LL (cyan). For each experimental repeat, a median bout duration within each 1-hour bin was calculated. Solid lines represent the means of medians. Shaded areas indicate the 95% confidence interval. **(C)** Lomb–Scargle periodograms of detrended bout duration under DD (red), LD (gray), and LL (cyan). Bout durations were binned in 1-hour intervals and summarized by per-fish medians. Dominant peak significance was assessed via permutation tests (p < 0.001 for all conditions). **(D)** Cosinor fits of detrended bout-duration time series under DD (red) and LL (cyan). Bout durations were binned into 1-hour intervals and summarized as per-fish medians. Single-component models were fit at the condition level with the period fixed to the dominant Lomb–Scargle period. Shaded regions indicate 95% confidence intervals. **(E)** IBI postural drift (upper) and bout rotation (lower) in DD plotted as a function of time over 5 days of recording. For each experimental repeat, a median value within each 1-hour bin was calculated. Solid lines represent the means of medians. Shaded areas indicate the 95% confidence interval. **(F)** Lomb–Scargle periodograms of detrended, 1-hour-binned per-fish median values for postural drift (top; permutation p < 0.001) and bout rotation (bottom; permutation p < 0.001). **(G)** Cosinor fits of detrended bout-rotation time series under DD. Bout rotation values were binned in 1-hour intervals and summarized as per-fish medians. Single-component models were fit with the period fixed to the dominant Lomb–Scargle period. Points show 1-hour binned medians; lines show Cosinor fits. n = 40080/135975/115146 DD/LD/LL bouts over 18/19/17 larvae. For calculation of IBI postural drift, only swim bouts with a recorded preceding bout were included. n = 23791 bouts over 18 larvae in E–G.

Larvae in constant darkness exhibited oscillations in bout duration (Figure 4B, red). Strikingly, larvae under LD exhibited a pronounced biphasic pattern with rapid transitions at light-dark switches (Figure 4B, black). In contrast, larvae in LL maintained short bout durations (Figure 4B, cyan), indicating a masking effect of light.

To characterize circadian periods, we detrended bout duration measurements to mitigate developmental effects and applied Lomb–Scargle periodogram analysis (Figure 4C). We observed strong dominant periods of 27.19 h, 24.13 h under DD and LD conditions, respectively, with powers of 0.19 and 0.61 (permutation p < 0.001). The longer-than-24-hour period in DD larvae suggests that we captured the free-running rhythm of larvae zebrafish [15]. Interestingly, the detected period was approximately 1 hour longer than previously reported in larvae zebrafish [15, 18]. In contrast to the robust rhythmic changes in bout duration observed under DD and LD, larvae under LL exhibited only week rhythmicity (period = 24.31 h, power = 0.03, permutation p < 0.001).

We further characterized circadian rhythmicity by applying Cosinor analysis to DD and LL datasets using the dominant Lomb–Scargle period (Figure 4D). DD larvae exhibited clear oscillations in bout duration, with longer bout duration during circadian night (Figure 4D, red; R^2^ = 0.191, amplitude = 16.4 ms). In contrast, LL larvae showed poor Cosinor fits (Figure 4D, cyan; R^2^ = 0.031, amplitude = 2.3 ms). The small estimated difference which is merely 4.5 ms between circadian day and night was smaller than the 6 ms temporal resolution limit of SAMPL [25].

Because longer bouts are associated with greater bout rotations that can compensate for larger postural drift (Figure 2), we next asked whether circadian oscillations in bout duration reflected corresponding rhythms in IBI postural drift and bout rotation. Across 5 days of behavioural recordings in DD larvae, IBI postural drift gradually decreased, whereas bout rotation gradually increased (Figure 4E). To isolate rhythmic components independent of long-term trends, we detrended these measurements and applied Lomb–Scargle periodogram analysis. We observed significant rhythmicity for both IBI postural drift and bout rotation, with similar periods close to 27 hours (Figure 4F, permutation p < 0.001 for both parameters; 26.96 h / 27.19 h periods for IBI drift / bout rotation). We further visualized oscillations in both parameters using Cosinor fits (Figure 4G). Notably, Cosinor fits for IBI postural drift and bout rotation showed comparable amplitudes (Table 4, 2.44 deg vs 2.26 deg), but approximately opposite phases (Table 4, 1.04 h vs 13.69 h). These results demonstrate that larvae adjust bout rotation magnitude in coordination with circadian fluctuations in postural drift.

**Table 4:** Circadian rhythm parameters for 2-bout latter bout duration, IBI rotation, and bout rotation. Refer to Figure 4. Lomb–Scargle and cosinor analyses were performed on detrended 1 h per-fish median values. Lomb–Scargle peaks were searched between 12 to 36 h periods, and empirical peak p-values were estimated with 1000 permutations. Cosinor fits used each condition’s dominant Lomb–Scargle period. Phase is the Cosinor acrophase in hours relative to the aligned experiment start, modulo the fitted period.

| Parameter | Unit | Cond. | Fish n | Lomb–Scargle |  |  | Cosinor |  |  |  |
| --- | --- | --- | --- | --- | --- | --- | --- | --- | --- | --- |
|  |  |  |  | Period (h) | Power | Perm. P | Amplitude | Phase (h) | R<sup>2</sup> | P-value |
| Bout duration | ms | DD | 12 | 27.190 | 0.181 | <0.001 | 17.345 | 15.690 | 0.184 | <0.001 |
|  |  | LD | 16 | 24.135 | 0.550 | <0.001 | — | — | — | — |
|  |  | LL | 12 | 24.690 | 0.033 | <0.001 | 2.542 | 0.147 | 0.033 | <0.001 |
| IBI postural drift | deg | DD | 12 | 26.962 | 0.072 | <0.001 | 2.437 | 1.035 | 0.074 | <0.001 |
|  |  | LD | 16 | 23.779 | 0.398 | <0.001 | — | — | — | — |
|  |  | LL | 12 | 25.271 | 0.088 | <0.001 | 2.352 | 10.986 | 0.089 | <0.001 |
| Bout rotation | deg | DD | 12 | 27.190 | 0.061 | <0.001 | 2.260 | 13.686 | 0.061 | <0.001 |
|  |  | LD | 16 | 23.779 | 0.371 | <0.001 | — | — | — | — |
|  |  | LL | 12 | 25.074 | 0.078 | <0.001 | 2.161 | 23.972 | 0.078 | <0.001 |

Together, these findings demonstrate that light exerts a masking effect that promotes the exploratory strategy, whereas the circadian clock regulates the magnitude of postural corrections in darkness. Our results disentangle the respective contributions of endogenous rhythms and environmental lighting to the regulation of locomotion in larval zebrafish.

## DISCUSSION

We defined two distinct locomotor strategies in larval zebrafish and demonstrated how lighting and circadian rhythms collectively shaped locomotion. Under light conditions, larvae used frequent, short swim bouts with increased directional variability that promoted horizontal exploration. In the absence of light, larvae performed long bouts with greater nose-up rotations that compensated for postural drifts accrued during inactivity. While light exerted a dominant masking effect that promoted the exploratory strategy, the circadian clock rhythmically modulated swim kinematics in darkness, with postural drift and corrective bout rotations exhibiting opposing circadian phases. Our results demonstrate how fish adjust swim strategies to achieve longer periods of inactivity while maintaining postural and depth control. This work elucidates how environmental cues and the internal clock shape locomotion strategies in a freely moving small vertebrate.

### Circadian and light modulation of balance and navigation

Circadian and lighting cues affect animal behaviour [1–8, 50]. Traditionally, rhythmic activities were quantified using measures such as wheel running, beam crossings, and total distance travelled – metrics that primarily served as circadian phase markers [16, 17, 51, 52]. Recent advancements in machine-learning-based video analysis have provided powerful tools for extracting posture and movement kinematics in freely moving animals [53, 54]. However, their intensive computational demands hinder their use in extended recordings. Conversely, approaches using automatic classification enabled tracking of rhythmic behaviour through multiple days [55, 56], but they often lack the resolution needed for detailed kinematic analysis in freely behaving animals. How changes in activity levels translate into kinematics that describe behavioural strategies remains largely unknown. We addressed these challenges by combining the simplicity of zebrafish behaviour with high-resolution, real-time pose estimation [25]. We achieved multi-day behavioural datasets by recording larvae that stochastically entered a 20 mm ×20 mm field of view in the centre of the arena [25]. This enabled us to define locomotor strategies in freely swimming fish and identify circadian regulation of postural control and locomotion.

Classic paradigms for zebrafish circadian studies use dim illumination ranging from 3 to 40 lux for the constant light condition [16, 17, 50, 57–60]. Larvae exposed to light intensities above this range were thought to be constantly active with a complete loss of circadian rhythm [60]. In our constant light condition, we used regular intensity light that was comparable to the photophase of the LD and observed a dominant effect of light on larval locomotor strategy, consistent with the classic “masking effect” [5–7]. Light-induced masking of behaviour has been reported across species, including mammals [8, 61, 62], birds [63], reptiles [64, 65], fish [18, 23, 24, 50, 60], and fruit flies [1, 66]. Nevertheless, mechanisms linking the modulation of sensory-motor circuits to the masking effect of light remain elusive. Unlike mammals, zebrafish can detect light directly via the pineal gland and in many other cell types [67–69]. Recent data indicate that both eyes and extra-retinal photosensitive tissues contribute to behavioural modulation in larval zebrafish [50, 70, 71]. Future studies combining loss-of-function approaches with deep phenotyping of light masking could elucidate the functional roles of different light-sensing pathways in locomotor behaviour.

Our results revealed rhythmic fluctuations in postural control kinematics in DD conditions, suggesting that balance behaviour could be modulated by the time of day. This finding adds to the limited evidence of circadian influences on balance [72, 73]. Larval zebrafish are an emerging model for studying vestibular function and underlying circuits [14, 26, 32, 74–76]. Recent work focusing on the zebrafish optomotor response has identified mechanisms through which circadian time modulates circuit dynamics and behaviour [77]. Future studies that integrate kinematic analysis of balance behaviour with measurement of vestibular circuit activity will be essential for uncovering how circadian rhythms shape vestibular processing and balance control.

Together, our work sets the stage for exploring neural mechanisms by which lighting conditions and the internal clock collectively modulate motor outputs for postural control and locomotion in a small, diurnal vertebrate.

### Compensatory body rotation implies short-term memory of vestibular signals

Zebrafish are diurnal, showing prolonged periods of inactivity between swim bouts in the dark or at night. We found that larvae compensate for postural drifts by swimming with significant nose-up rotations. This strategy enables longer inactive periods while minimizing unfavourable postures. Unexpectedly, the amount of compensatory rotation appeared to be associated with the amount of passive postural drift rather than with the duration of the inactive phase. To achieve this, larvae might need to remember either their posture following a swim bout or postural drift during the inactive phase. These findings suggest the existence of a short-term memory of vestibular signals on the pitch axis.

Classic heading direction cells integrate angular velocity over time to encode directional and positional signals in the horizontal plane [78–82]. In complete darkness, vertebrates rely on vestibular input to maintain directional tuning when external reference points are unavailable [78]. Unlike horizontal rotation, where only angular velocity can be detected, tilt angles in vertical planes (pitch and roll) can be inferred from gravitational signals detected by the otolith organs in the inner ear [83]. In larval zebrafish, the semicircular canals remain non-functional during the period we studied here [84]. Nevertheless, the inner-ear otoliths enable sensation of gravity and linear acceleration [75, 85]. Evolutionarily conserved brainstem vestibular circuits transform otolithic input into motor outputs that control eye movements, stabilize posture, and maintain swim directions [26, 32, 45, 86, 87]. Our results demonstrate that larvae rely on otolithic signals to restore heading on the pitch axis following prolonged inactivity. Future studies may identify the neural substrates responsible for the storage of vestibular signals that enable postural compensation.

By characterizing the compensatory body rotation, our work lays the foundation for investigating neural mechanisms of vestibular processing using larval zebrafish.

### Limitations

One statistical limitation of this study is the relatively small number of independent experimental repeats (n = 5), which reduces statistical power for repeat-level comparisons. Consequently, some parameters exhibited large effect sizes but only marginal p-values. To assess the robustness of these results, we performed complementary bout-level analyses, which leverages substantially greater sample sizes at the level of individual swim bouts. These analyses yielded consistent estimates across parameters and are summarized in Tables 1 and 3.

The behavioural data acquired in this study are subject to several limitations arising from design choices and technical compromises of our apparatus.

First, the size of our behaviour chamber raises questions about the ecological relevance of exploration in this context. Compared to previous studies that assessed exploratory swimming in 35 mm dishes holding 5 mL of water [49, 88], our setup contained 5 to 7 larvae swimming in 30 mL of water measured approximately 50.8 mm (L) × 50 mm (H) × 12.7 mm (W) [25]. Given that the volume of a 7 dpf larva, approximately 4 mm in length and 0.6 mm in width, can be estimated at around 0.4 to 0.8 mm^3^ [89], our setup is roughly equivalent to humans diving in an Olympic-sized swimming pool. A larva swimming continuously in the same direction would take 25 to 50 seconds to cross the chamber. Thus, we reason that we measured kinematic parameters of larvae exploring the environment. It is plausible that by introducing novel environments and points of interest, future studies will reveal more sophisticated exploratory strategies and enhance ecological validity.

Second, to achieve high-resolution imaging over 48 hours, we recorded larvae that stochastically entered a 20 mm × 20 mm field of view in the centre of the arena [25]. While this approach enabled detailed analyses of locomotor kinematics of individual fish, it was less suitable for measuring the free-running period than for capturing the global activity of all larvae. Moreover, although we observed fast adaptation of locomotor strategies to light transitions, we could not determine how quickly larvae adjusted their swim strategy during the light transition. Previous work revealed distinct acute responses to loss of illumination within 3 minutes [71, 90], suggesting that our recording likely missed acute responses but captured longer-term effects of changes in lighting. In addition, due to the limited field of view, we could not determine the percentage of time larvae swimming in the water column vs. staying at the bottom. Although they, on average, adopted 13° climbs in the dark – potentially compensating for sinking between bouts – larvae likely also rest for extended periods on the bottom of the arena. Future studies using a deeper arena with global activity tracking would help elucidate the full range of resting and sleep behaviours in fish.

Lastly, because the white-light LED strips and their usage varied across apparatuses, recordings under light conditions were obtained at illuminance levels ranging from 50 to 150 lux. The corresponding unweighted broadband irradiance was quantified and found to span 0.61 to 3.91 W/m^2^. We compared behaviour parameters across all boxes and did not detect systematic relationships between irradiance and measurements. Future studies with precisely controlled illumination will reveal how light intensity influences postural control and navigation in larval zebrafish.

### Conclusion

We demonstrated that lighting and circadian cues shape locomotor strategies in larval zebrafish, prioritizing postural control in the dark and environmental exploration in the light. Our findings highlight how the interplay of external and internal cues governs balance and navigation in a freely moving small vertebrate.

## ACKNOWLEDGMENTS

Research was supported by the National Institute on Deafness and Communication Disorders of the National Institutes of Health under award numbers R01DC017489 (DS), the National Institute of Neurological Disorders and Stroke under award number R33NS125280 (DS), and the Rainwater Charitable Foundation (YZ). The authors dedicate this manuscript to the memory of Prof. Michael Menaker.

## AUTHOR CONTRIBUTIONS

Conceptualization: JL; Methodology: YZ, JL, AK; Investigation: JL, YZ, AK, SND, HG, NM; Visualization: JL, YZ; Writing: JL, YZ; Editing: YZ, DS; Funding Acquisition: DS, YZ; Supervision: YZ, DS.

---

**Figure S1:**
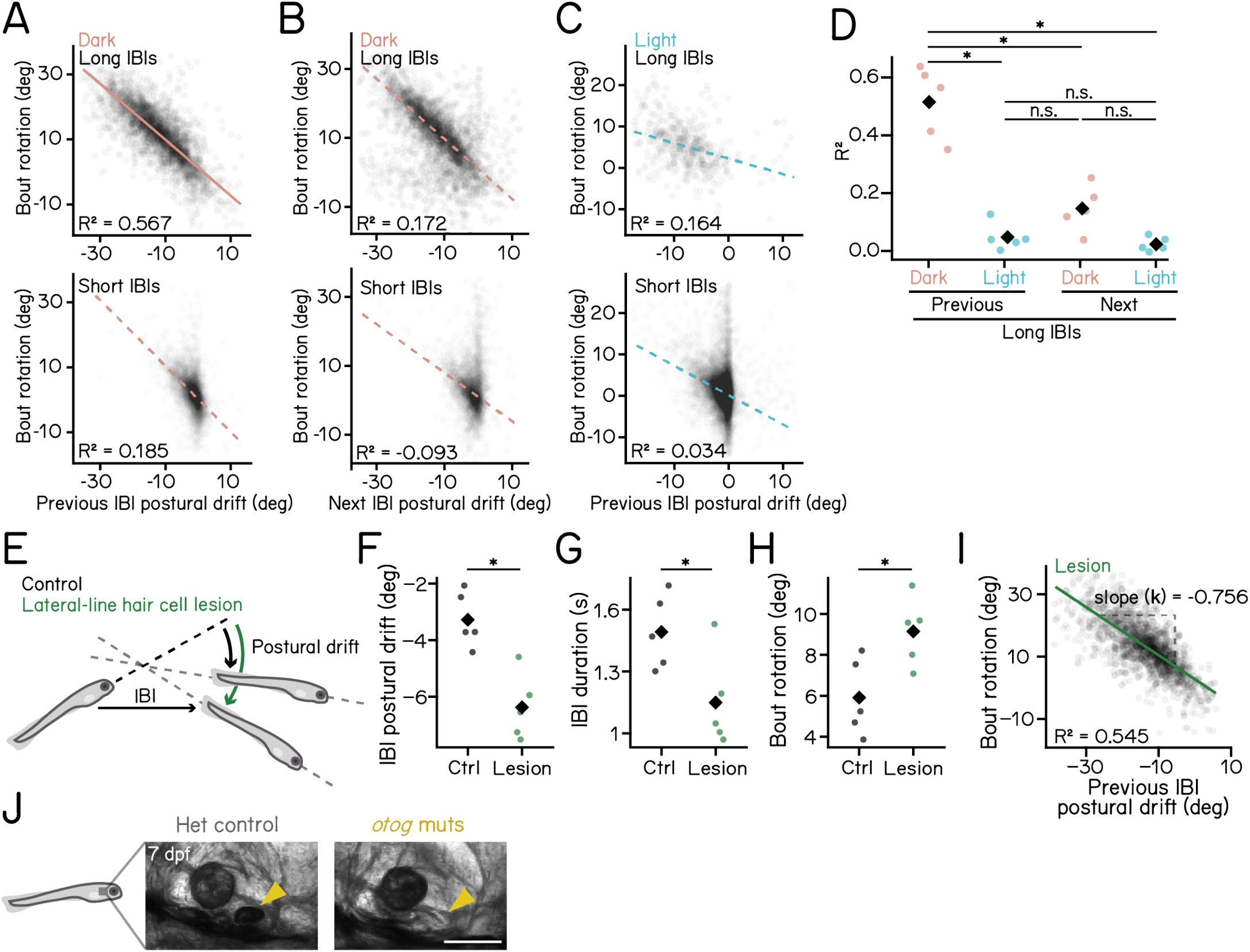
Bout rotations following long swim intervals correlate with IBI postural drifts. Refer to Figure 2. **(A)** Scatter plot of bout rotation vs. postural drift during the previous IBI in the dark. Bouts were categorized into long and short IBIs based on the duration of preceding IBIs. Red lines represent the robust bi-square regression fit. Slope of the best-fit line for long IBIs: −0.860, *R*^2^ = 0.567; slope for short IBIs: −0.986, *R*^2^ = 0.185. **(B)** Scatter plot of bout rotation vs. postural drift during the following IBI in the dark. Bouts were categorized into long and short IBIs based on the duration of following IBIs. Red lines represent the robust bi-square regression fit. Slope of the best-fit line for long IBIs: −0.874, *R*^2^ = 0.172; slope for short IBIs: −0.712, *R*^2^ = −0.093. **(C)** Scatter plot of bout rotation vs. postural drift during the previous IBI in the light. Bouts were categorized into long and short IBIs based on the duration of preceding IBIs. Cyan lines represent the robust bi-square regression fit. Slope of the best-fit line for long IBIs: −0.360, *R*^2^ = 0.164; short IBIs: −0.753, and *R*^2^ = 0.034. **(D)** Comparison of *R*^2^ values for regression fits of bout rotation against IBI postural drift (*: adjusted p < 0.001, two-way ANOVA with post-hoc Tukey HSD tests). **(E)** Schematics illustrating effects of lateral-line hair cell lesions on IBI postural drifts. **(F–H)** Comparisons of bout kinematics between lateral-line lesioned larvae and sham controls: IBI rotation (p = 1.882e-03, Cohen’s d = 2.876, t-test) **(F)**, IBI duration (p = 2.992e-02, Cohen’s d = 1.667, t-test) **(G)**, and bout rotation (p = 2.028e-02, Cohen’s d = 1.826, t-test) **(H)**. Median values for each experimental repeat are plotted as dots. Diamonds indicate group means. **(I)**Scatter plot of bout rotation vs. drift during the previous IBI following lesions. Only bouts with long preceding IBI were plotted. The green line represents the robust bi-square regression fit. Slope of the best-fit line for long IBIs: −0.756, coefficient of determination *R*^2^ = 0.545. **(J)** *otog* mutants lack the utricular otolith (arrowheads) at 7 dpf. Scale bar: 100 µm.Middle bouts from 3-bout sequences were selected for IBI postural drift computation. n = 3143/34285 day-time bouts from 105/98 fish over 5 experimental repeats for wild type DD/LD. For (F–H), n = 25008/8982 day-time bouts from 114/114 fish over 5 experimental repeats for lateral-line hair cell control/lesions. For (I), n = 2994 middle bouts from 3-bout. See also Table 2.

## REFERENCES

[1] Bünning E. Zur kenntnis der endonomen tagesrhythmik bei insekten und bei pflanzen. Berichte der Deutschen Botanischen Gesellschaft, 53(7):594623, October 1935.

[2] C. S. Pittendrigh. On temporal organization in living systems. Harvey Lectures, 56:93–125, 1960. Published 1960–1961.

[3] N. Mrosovsky, S. G. Reebs, G. I. Honrado, and P. A. Salmon. Behavioural entrainment of circadian rhythms. Experientia, 45(8):696702, August 1989.

[4] Tara A. LeGates, Diego C. Fernandez, and Samer Hattar. Light as a central modulator of circadian rhythms, sleep and affect. Nature Reviews Neuroscience, 15(7):443454, June 2014.

[5] J. Aschoff. Exogenous and endogenous components in circadian rhythms. Cold Spring Harbor Symposia on Quantitative Biology, 25(0):1128, January 1960.

[6] M. D. Marques and J. M. Waterhouse. Masking and the evolution of circadian rhythmicity. Chronobiology International, 11(3):146155, January 1994.

[7] N. Mrosovsky. Masking: History, definitions, and measurement. Chronobiology International, 16(4):415429, January 1999.

[8] Lily Yan, Laura Smale, and Antonio A. Nunez. Circadian and photic modulation of daily rhythms in diurnal mammals. European Journal of Neuroscience, 51(1):551566, October 2018.

[9] Serge Daan. Adaptive Daily Strategies in Behavior, page 275298. Springer US, 1981.

[10] C S Pittendrigh. Temporal organization: Reflections of a darwinian clock-watcher. Annual Review of Physiology, 55(1):1754, October 1993.

[11] Vincent van der Vinne, Jenke A. Gorter, Sjaak J. Riede, and Roelof A. Hut. Diurnality as an energy-saving strategy: energetic consequences of temporal niche switching in small mammals. Journal of Experimental Biology, 218(16):25852593, August 2015.

[12] Yu. G. Aleyev. Nekton. Springer Netherlands, 1977.

[13] Martha W Bagnall and David Schoppik. Development of vestibular behaviors in zebrafish. Current Opinion in Neurobiology, 53:8389, December 2018.

[14] David E. Ehrlich and David Schoppik. Control of movement initiation underlies the development of balance. Current Biology, 27(3):334344, February 2017.

[15] Gregory M. Cahill, Mark W. Hurd, and Matthew M. Batchelor. Circadian rhythmicity in the locomotor activity of larval zebrafish. NeuroReport, 9(15):34453449, October 1998.

[16] M Hurd, J Debruyne, M Straume, and G Cahill. Circadian rhythms of locomotor activity in zebrafish. Physiology & Behavior, 65(3):465472, October 1998.

[17] Gregory M. Cahill. Clock mechanisms in zebrafish. Cell and Tissue Research, 309(1):2734, July 2002.

[18] Mark W. Hurd and Gregory M. Cahill. Entraining signals initiate behavioral circadian rhythmicity in larval zebrafish. Journal of Biological Rhythms, 17(4):307314, August 2002.

[19] Harold A. Burgess and Michael Granato. Modulation of locomotor activity in larval zebrafish during light adaptation. Journal of Experimental Biology, 210(14):25262539, July 2007.

[20] David A. Prober, Jason Rihel, Anthony A. Onah, Rou-Jia Sung, and Alexander F. Schier. Hypocretin/orexin overexpression induces an insomnia-like phenotype in zebrafish. The Journal of Neuroscience, 26(51):1340013410, December 2006.

[21] Grigorios Oikonomou and David A Prober. Attacking sleep from a new angle: contributions from zebrafish. Current Opinion in Neurobiology, 44:8088, June 2017.

[22] Louis C. Leung, Gordon X. Wang, Romain Madelaine, Gemini Skariah, Koichi Kawakami, Karl Deisseroth, Alexander E. Urban, and Philippe Mourrain. Neural signatures of sleep in zebrafish. Nature, 571(7764):198204, July 2019.

[23] Qian Lin and Suresh Jesuthasan. Masking of a circadian behavior in larval zebrafish involves the thalamo-habenula pathway. Scientific Reports, 7(1), June 2017.

[24] Viacheslav V. Krylov, Evgeny I. Izvekov, Vera V. Pavlova, Natalia A. Pankova, and Elena A. Osipova. Circadian rhythms in zebrafish (danio rerio) behaviour and the sources of their variability. Biological Reviews, 96(3):785797, December 2020.

[25] Yunlu Zhu, Franziska Auer, Hannah Gelnaw, Samantha N. Davis, Kyla R. Hamling, Christina E. May, Hassan Ahamed, Niels Ringstad, Katherine I. Nagel, and David Schoppik. Sampl is a high-throughput solution to study unconstrained vertical behavior in small animals. Cell Reports, 42(6):112573, June 2023.

[26] Yunlu Zhu, Hannah Gelnaw, Franziska Auer, Kyla R. Hamling, David E. Ehrlich, and David Schoppik. Evolutionarily conserved brainstem architecture enables gravity-guided vertical navigation. PLOS Biology, 22(11):e3002902, November 2024.

[27] Benjamin W. Lindsey, Frank M. Smith, and Roger P. Croll. From inflation to flotation: Contribution of the swimbladder to whole-body density and swimming depth during development of the zebrafish (danio rerio).Zebrafish, 7(1):8596, March 2010.

[28] Alexandra Venuto, Stacey Thibodeau-Beganny, Josef G. Trapani, and Timothy Erickson. A sensation for inflation: initial swim bladder inflation in larval zebrafish is mediated by the mechanosensory lateral line. Journal of Experimental Biology, 226(11), June 2023.

[29] Takeshi Yoshimatsu, Cornelius Schröder, Noora E. Nevala, Philipp Berens, and Tom Baden. Fovea-like photoreceptor specializations underlie single uv cone driven prey-capture behavior in zebrafish. Neuron, 107(2):320–337.e6, July 2020.

[30] António M. Fernandes, Kandice Fero, Aristides B. Arrenberg, Sadie A. Bergeron, Wolfgang Driever, and Harold A. Burgess. Deep brain photoreceptors control light-seeking behavior in zebrafish larvae. Current Biology, 22(21):20422047, November 2012.

[31] Benjamin H. Bishop, Nathan Spence-Chorman, and Ethan Gahtan. Three-dimensional motion tracking reveals a diving component to visual and auditory escape swims in zebrafish larvae. Journal of Experimental Biology, January 2016.

[32] Kyla R. Hamling, Katherine Harmon, Yukiko Kimura, Shin-ichi Higashijima, and David Schoppik. The vestibulospinal nucleus is a locus of balance development. The Journal of Neuroscience, 44(30):e2315232024, May 2024.

[33] Samantha N. Davis, Yunlu Zhu, and David Schoppik. Larval zebrafish maintain elevation with multisensory control of posture and locomotion. bioRxiv, January 2024.

[34] Catherine O. Fritz, Peter E. Morris, and Jennifer J. Richler. Effect size estimates: Current use, calculations, and interpretation. Journal of Experimental Psychology: General, 141(1):218, 2012.

[35] DAVID W. Scott. On optimal and data-based histograms. Biometrika, 66(3):605610, 1979.

[36] Rainer Voigt, Alexander G. Carton, and John C. Montgomery. Responses of anterior lateral line afferent neurones to water flow. Journal of Experimental Biology, 203(16):24952502, August 2000.

[37] M.J. McHenry, K.E. Feitl, J.A. Strother, and W.J. Van Trump. Larval zebrafish rapidly sense the water flow of a predators strike. Biology Letters, 5(4):477479, March 2009.

[38] William J. Stewart, Gilberto S. Cardenas, and Matthew J. McHenry. Zebrafish larvae evade predators by sensing water flow. Journal of Experimental Biology, 216(3):388398, February 2013.

[39] Andres Carrillo and Matthew J. McHenry. Zebrafish learn to forage in the dark. Journal of Experimental Biology, 219(4):582589, February 2016.

[40] Pablo Oteiza, Iris Odstrcil, George Lauder, Ruben Portugues, and Florian Engert. A novel mechanism for mechanosensory-based rheotaxis in larval zebrafish. Nature, 547(7664):445448, 2017.

[41] T. J. Pitcher, B. L. Partridge, and C. S. Wardle. A blind fish can school. Science, 194(4268):963965, November 1976.

[42] Sheryl Coombs and Horst Bleckmann. The Gems of the Past: A Brief History of Lateral Line Research in the Context of the Hearing Sciences, page 116. Springer New York, 2013.

[43] Tanya T. Whitfield, Michael Granato, Fredericus J. M. van Eeden, Ursula Schach, Michael Brand, Makoto Furutani-Seiki, Pascal Haffter, Matthias Hammerschmidt, Carl-Philipp Heisenberg, Yun-Jin Jiang, Donald A. Kane, Robert N. Kelsh, Mary C. Mullins, Jörg Odenthal, and Christiane Nüsslein-Volhard. Mutations affecting development of the zebrafish inner ear and lateral line. Development, 123(1):241254, December 1996.

[44] Weike Mo, Fangyi Chen, Alex Nechiporuk, and Teresa Nicolson. Quantification of vestibular-induced eye movements in zebrafish larvae. BMC Neuroscience, 11(1), September 2010.

[45] Isaac H. Bianco, Leung-Hang Ma, David Schoppik, Drew N. Robson, Michael B. Orger, James C. Beck, Jennifer M. Li, Alexander F. Schier, Florian Engert, and Robert Baker. The tangential nucleus controls a gravito-inertial vestibulo-ocular reflex. Current Biology, 22(14):12851295, July 2012.

[46] Richard Roberts, Jeffrey Elsner, and Martha W. Bagnall. Delayed otolith development does not impair vestibular circuit formation in zebrafish. Journal of the Association for Research in Otolaryngology, 18(3):415425, March 2017.

[47] Bruce B. Riley and Stephen J. Moorman. Development of utricular otoliths, but not saccular otoliths, is necessary for vestibular function and survival in zebrafish. Journal of Neurobiology, 43(4):329337, 2000.

[48] R.C. MacPhail, J. Brooks, D.L. Hunter, B. Padnos, T.D. Irons, and S. Padilla. Locomotion in larval zebrafish: Influence of time of day, lighting and ethanol. NeuroToxicology, 30(1):5258, January 2009.

[49] Gokul Rajan, Julie Lafaye, Giulia Faini, Martin Carbo-Tano, Karine Duroure, Dimitrii Tanese, Thomas Panier, Raphaël Candelier, Jörg Henninger, Ralf Britz, Benjamin Judkewitz, Christoph Gebhardt, Valentina Emiliani, Georges Debregeas, Claire Wyart, and Filippo Del Bene. Evolutionary divergence of locomotion in two related vertebrate species. Cell Reports, 38(13):110585, March 2022.

[50] Clair Chaigne, Dora Sapède, Xavier Cousin, Laurent Sanchou, Patrick Blader, and Elise Cau. Contribution of the eye and of opn4xa function to circadian photoentrainment in the diurnal zebrafish. PLOS Genetics, 20(2):e1011172, February 2024.

[51] Joseph S. Takahashi and Martin Zatz. Regulation of circadian rhythmicity. Science, 217(4565):11041111, September 1982.

[52] Robert Lee, Amaris Tapia, Sevag Kaladchibachi, Michael A. Grandner, and Fabian-Xosé Fernandez. Meta-analysis of light and circadian timekeeping in rodents. Neuroscience &Biobehavioral Reviews, 123:215229, April 2021.

[53] Yujia Hu, Carrie R. Ferrario, Alexander D. Maitland, Rita B. Ionides, Anjesh Ghimire, Brendon Watson, Kenichi Iwasaki, Hope White, Yitao Xi, Jie Zhou, and Bing Ye. Labgym: Quantification of user-defined animal behaviors using learning-based holistic assessment. Cell Reports Methods, 3(3):100415, March 2023.

[54] Jialin Ye, Yang Xu, Kang Huang, Xinyu Wang, Liping Wang, and Feng Wang. Hierarchical behavioral analysis framework as a platform for standardized quantitative identification of behaviors. Cell Reports, 44(2):115239, February 2025.

[55] Logan J. Perry, Gavin E. Ratcliff, Arthur Mayo, Blanca E. Perez, Larissa Rays Wahba, K.L. Nikhil, William C. Lenzen, Yangyuan Li, Jordan Mar, Isabella Farhy-Tselnicker, Wanhe Li, and Jeff R. Jones. A circadian behavioral analysis suite for real-time classification of daily rhythms in complex behaviors. Cell Reports Methods, 5(5):101050, May 2025.

[56] Nikolas A. Francis, Kayla Bohlke, and Patrick O. Kanold. Automated behavioral experiments in mice reveal periodic cycles of task engagement within circadian rhythms. eneuro, pages ENEURO.0121–19.2019, September 2019.

[57] José F. LópezOlmeda, Juan A. Madrid, and Francisco J. SánchezVázquez. Light and temperature cycles as zeitgebers of zebrafish (danio rerio) circadian activity rhythms. Chronobiology International, 23(3):537550, January 2006.

[58] Lior Appelbaum, Gordon X. Wang, Geraldine S. Maro, Rotem Mori, Adi Tovin, Wilfredo Marin, Tohei Yokogawa, Koichi Kawakami, Stephen J. Smith, Yoav Gothilf, Emmanuel Mignot, and Philippe Mourrain. Sleepwake regulation and hypocretinmelatonin interaction in zebrafish. Proceedings of the National Academy of Sciences, 106(51):2194221947, December 2009.

[59] Adi Tovin, Shahar Alon, Zohar Ben-Moshe, Philipp Mracek, Gad Vatine, Nicholas S. Foulkes, Jasmine Jacob-Hirsch, Gideon Rechavi, Reiko Toyama, Steven L. Coon, David C. Klein, Eli Eisenberg, and Yoav Gothilf. Systematic identification of rhythmic genes reveals camk1gb as a new element in the circadian clockwork. PLoS Genetics, 8(12):e1003116, December 2012.

[60] Idan Elbaz, Nicholas S. Foulkes, Yoav Gothilf, and Lior Appelbaum. Circadian clocks, rhythmic synaptic plasticity and the sleep-wake cycle in zebrafish. Frontiers in Neural Circuits, 7, 2013.

[61] P. H. Gander and M. C. Moore-Ede. Light-dark masking of circadian temperature and activity rhythms in squirrel monkeys. American Journal of Physiology-Regulatory, Integrative and Comparative Physiology, 245(6):R927R934, December 1983.

[62] Noga Kronfeld-Schor, Davide Dominoni, Horacio de la Iglesia, Oren Levy, Erik D. Herzog, Tamar Dayan, and Charlotte Helfrich-Forster. Chronobiology by moonlight. Proceedings of the Royal Society B: Biological Sciences, 280(1765):20123088, August 2013.

[63] Jürgen Aschoff and Christiana von Goetz. Masking of circadian activity rhythms in canaries by light and dark. Journal of Biological Rhythms, 4(1):2938, March 1989.

[64] Herbert Underwood and Michael Menaker. Extraretinal light perception: Entrainment of the biological clock controlling lizard locomotor activity. Science, 170(3954):190193, October 1970.

[65] Cristiano Bertolucci, Valeria Anna Sovrano, Maria Chiara Magnone, and Augusto Foà. Role of suprachiasmatic nuclei in circadian and light-entrained behavioral rhythms of lizards. American Journal of Physiology-Regulatory, Integrative and Comparative Physiology, 279(6):R2121R2131, December 2000.

[66] Dirk Rieger, Ralf Stanewsky, and Charlotte Helfrich-Förster. Cryptochrome, compound eyes, hofbauer-buchner eyelets, and ocelli play different roles in the entrainment and masking pathway of the locomotor activity rhythm in the fruit fly drosophila melanogaster. Journal of Biological Rhythms, 18(5):377391, October 2003.

[67] David Whitmore, Nicholas S. Foulkes, and Paolo Sassone-Corsi. Light acts directly on organs and cells in culture to set the vertebrate circadian clock. Nature, 404(6773):8791, March 2000.

[68] Amanda-Jayne F. Carr and David Whitmore. Imaging of single light-responsive clock cells reveals fluctuating free-running periods. Nature Cell Biology, 7(3):319321, March 2005.

[69] Helen A. Moore and David Whitmore. Circadian rhythmicity and light sensitivity of the zebrafish brain. PLoS ONE, 9(1):e86176, January 2014.

[70] Zohar Ben-Moshe Livne, Shahar Alon, Daniela Vallone, Yared Bayleyen, Adi Tovin, Inbal Shainer, Laura G. Nisembaum, Idit Aviram, Sima Smadja-Storz, Michael Fuentes, Jack Falcón, Eli Eisenberg, David C. Klein, Harold A. Burgess, Nicholas S. Foulkes, and Yoav Gothilf. Genetically blocking the zebrafish pineal clock affects circadian behavior. PLOS Genetics, 12(11):e1006445, November 2016.

[71] Eric J. Horstick, Yared Bayleyen, Jennifer L. Sinclair, and Harold A. Burgess. Search strategy is regulated by somatostatin signaling and deep brain photoreceptors in zebrafish. BMC Biology, 15(1), January 2017.

[72] Tristan Martin, Amira Zouabi, Florane Pasquier, Pierre Denise, Antoine Gauthier, and Gaëlle Quarck. Twenty-four-hour variation of vestibular function in young and elderly adults. Chronobiology International, 38(1):90102, December 2020.

[73] Alex I. Halpern, Jamie A.F. Jansen, Nir Giladi, Anat Mirelman, and Jeffrey M. Hausdorff. Does time of day influence postural control and gait? a review of the literature. Gait & Posture, 92:153166, February 2022.

[74] David E Ehrlich and David Schoppik. A primal role for the vestibular sense in the development of coordinated locomotion. eLife, 8, October 2019.

[75] Zhikai Liu, David G. C. Hildebrand, Joshua L. Morgan, Yizhen Jia, Nicholas Slimmon, and Martha W. Bagnall. Organization of the gravity-sensing system in zebrafish. Nature Communications, 13(1), August 2022.

[76] Kyla R. Hamling, Katherine Harmon, and David Schoppik. The nature and origin of synaptic inputs to vestibulospinal neurons in the larval zebrafish. eneuro, 10(6):ENEURO.0090–23.2023, June 2023.

[77] Patrício Simões, José Moya-Díaz, and Leon Lagnado. Quantifying the link between retinal performance and the optomotor response. Current Biology, July 2025.

[78] Jeffrey S. Taube. The head direction signal: Origins and sensory-motor integration. Annual Review of Neuroscience, 30(1):181207, July 2007.

[79] Edvard I. Moser, Emilio Kropff, and May-Britt Moser. Place cells, grid cells, and the brains spatial representation system. Annual Review of Neuroscience, 31(1):6989, July 2008.

[80] Kayvon Daie, Markă S. Goldman, and Emreă R.F. Aksay. Spatial patterns of persistent neural activity vary with the behavioral context of short-term memory. Neuron, 85(4):847860, February 2015.

[81] Daniel Turner-Evans, Stephanie Wegener, Hervé Rouault Romain Franconville, Tanya Wolff, Johannes D Seelig, Shaul Druckmann, and Vivek Jayaraman. Angular velocity integration in a fly heading circuit. eLife, 6, May 2017.

[82] Luigi Petrucco, Hagar Lavian, You Kure Wu, Fabian Svara, Vilim tih, and Ruben Portugues. Neural dynamics and architecture of the heading direction circuit in zebrafish. Nature Neuroscience, 26(5):765773, April 2023.

[83] D. E. Angelaki and B. J. Hess. Three-dimensional organization of otolith-ocular reflexes in rhesus monkeys. i. linear acceleration responses during off-vertical axis rotation. Journal of Neurophysiology, 75(6):24052424, June 1996.

[84] Selina Baeza-Loya and David W. Raible. Vestibular physiology and function in zebrafish. Frontiers in Cell and Developmental Biology, 11, April 2023.

[85] Kacey Mackowetzky, Kevin H. Yoon, Emily J. Mackowetzky, and Andrew J. Waskiewicz. Development and evolution of the vestibular apparatuses of the inner ear. Journal of Anatomy, 239(4):801828, May 2021.

[86] David Schoppik, Isaac H. Bianco, David A. Prober, Adam D. Douglass, Drew N. Robson, Jennifer M.B. Li, Joel S.F. Greenwood, Edward Soucy, Florian Engert, and Alexander F. Schier. Gaze-stabilizing central vestibular neurons project asymmetrically to extraocular motoneuron pools. The Journal of Neuroscience, 37(47):1135311365, September 2017.

[87] Celine Bellegarda, Franziska Auer, and David Schoppik. Zebrafish as a model to understand extraocular motor neuron diversity. Current Opinion in Neurobiology, 90:102964, February 2025.

[88] William A. Haney, Bushra Moussaoui, and James A. Strother. Prolonged exposure to stressors suppresses exploratory behavior in zebrafish larvae. Journal of Experimental Biology, January 2020.

[89] Yuanhao Guo, Wouter J. Veneman, Herman P. Spaink, and Fons J. Verbeek. Three-dimensional reconstruction and measurements of zebrafish larvae from high-throughput axial-view in vivo imaging. Biomedical Optics Express, 8(5):2611, April 2017.

[90] Matthew R. Waalkes, Maegan Leathery, Madeline Peck, Allison Barr, Alexander Cunill, John Hageter, and Eric J. Horstick. Light wavelength modulates search behavior performance in zebrafish. Scientific Reports, 14(1), July 2024.

